# Polishing *de novo* nanopore assemblies of bacteria and eukaryotes with FMLRC2

**DOI:** 10.1101/2022.07.22.501182

**Authors:** Q. Charles Mak, Ryan R. Wick, James Matthew Holt, Jeremy R. Wang

## Abstract

As accuracy and throughput of nanopore sequencing improves, it is increasingly common to perform long-read-first *de novo* genome assemblies followed by polishing with accurate short reads (Kim *et al*. 2021). We briefly introduce FMLRC2, the successor to the original FM-index Long Read Corrector (FMLRC), and illustrate its performance as a fast and accurate *de novo* assembly polisher for both microbial and eukaryotic genomes.

## Introduction

Long-read, third-generation sequencing technologies including Oxford Nanopore Technologies (ONT) and Pacific Biosciences (PacBio) are increasingly the workhorse and backbone of *de novo* genome assemblies (Amarasinghe et al., 2020; Pollard et al., 2018). Median read lengths from 10s to 100s of kilobases are routinely achieved (Michael et al., 2018; Shi et al., 2016), with which modern assemblers produce much more contiguous and complete *de novo* assemblies than those from short-read next-generation sequencing (NGS) alone (Amarasinghe et al., 2020; Pollard et al., 2018). Despite continuous improvement in nucleotide-level accuracy of long-read sequencing, residual errors - both single-nucleotide mismatches and short insertions and deletions (indels) - still significantly exceed short-read sequencing-by-synthesis technologies (Amarasinghe et al., 2020; Pollard et al., 2018).

Residual consensus errors in long-read assemblies are dominated by indels which hinder gene annotation (Watson & Warr, 2019). A “hybrid” assembly approach is commonly taken to maximize assembly accuracy and contiguity by first producing a draft assembly from long-read sequences, followed by polishing with accurate short reads (Jain et al., 2018).

FM-index Long Read Corrector (FMLRC; Wang et al., 2018) is a hybrid error-correction method that employs an FM-indexed BWT built from accurate reads to dynamically reassemble erroneous subregions of error-prone long sequences. FMLRC has proven a consistently accurate and efficient method for correcting sequencing errors in raw long reads (Zhang et al., 2020; Fu et al., 2019). FMLRC2 produces largely identical results to FMLRC albeit with significantly improved speed and stability. In addition to its proven utility for raw error correction, we demonstrate the effectiveness of FMLRC2 as a *de novo* assembly polisher for diverse prokaryotic and eukaryotic genomes. FMLRC2 consistently outperforms extant genome polishing tools in minimizing residual assembly errors (mismatches and indels) and computational requirements.

## Methods

### FMLRC2

FMLRC2 represents a reimplementation of the original FMLRC from C++ to Rust, without major changes to the underlying algorithm. In benchmark tests, it is about 50% faster than FMLRC. FMLRC2 is open source and publicly available at https://github.com/HudsonAlpha/fmlrc2.

### Datasets

We evaluated FMLRC2 against *de novo* assemblies from 24 microbial and six eukaryotic datasets. Bacterial datasets include four independent datasets from each of six bacterial isolates (*A. baumannii* J9, *C. koseri* MINF_9D, *E. kobei* MSB1_1B, Haemophilus M1C132_1, *K. oxytoca* MSB1_2C and *K. variicola* INF345) (Wick et al., 2021a). Long-read-only assemblies for each were performed using Trycycler v0.5.0 (Wick et al., 2021b) and Medaka v1.4.3 (Wright and Wykes). We additionally used two publicly available sets of ONT sequence data from each of three well-established model eukaryotic organisms: *Saccharomyces cerevisiae* (S288C), *Arabidopsis thaliana* (Columbia; TAIR10.1), and *Drosophila melanogaster* (ISO-1) [Table 1]. Experimental sequencing datasets (ONT and Illumina/BGI) were obtained from NCBI (Table 2). We simulated ONT data and the corresponding paired-end Illumina dataset using Badread v0.2.0 (Wick, 2019) and ART v2016-06-05 (Huang et al., 2011) as previously described (Wick and Holt, 2022). Briefly, short reads were simulated using ART with HiSeqX TruSeq preset, to 100X effective sequencing depth, 150 bp read length, 400±50(sd) bp mean fragment. Long reads were simulated using Badread with parameters “--length 20000,12000 --identity 90,98,4”. Basic statistics and coverage of simulated data are described in Table 3. FastQC v0.11.9 (Andrew, 2010) was used to check for quality issues among experimental short-read datasets. Where necessary, fastp v0.23.2 (Chen et al., 2018) was used to clean the short read datasets and remove Ns prior to downstream polishing. Long-read-only assemblies were generated for each of the nine experimental and simulated eukaryotic ONT datasets using Flye v2.8.1 (Kolmogorov et al., 2019) with default parameters.

**Table 1.** Eukaryotic model organisms used for assembly polishing evaluation

| Species | Strain/Genome | Refseq accession | Genome size |
| --- | --- | --- | --- |
| *Saccharomyces cerevisiae* | S288C | GCF_000146045.2 | 12.16 Mbp |
| *Arabidopsis thaliana* | Columbia (TAIR10.1) | GCF_000001735.4 | 119.67 Mbp |
| *Drosophila melanogaster* | ISO-1 | GCF_000001215.4 | 143.73 Mbp |

**Table 2:**
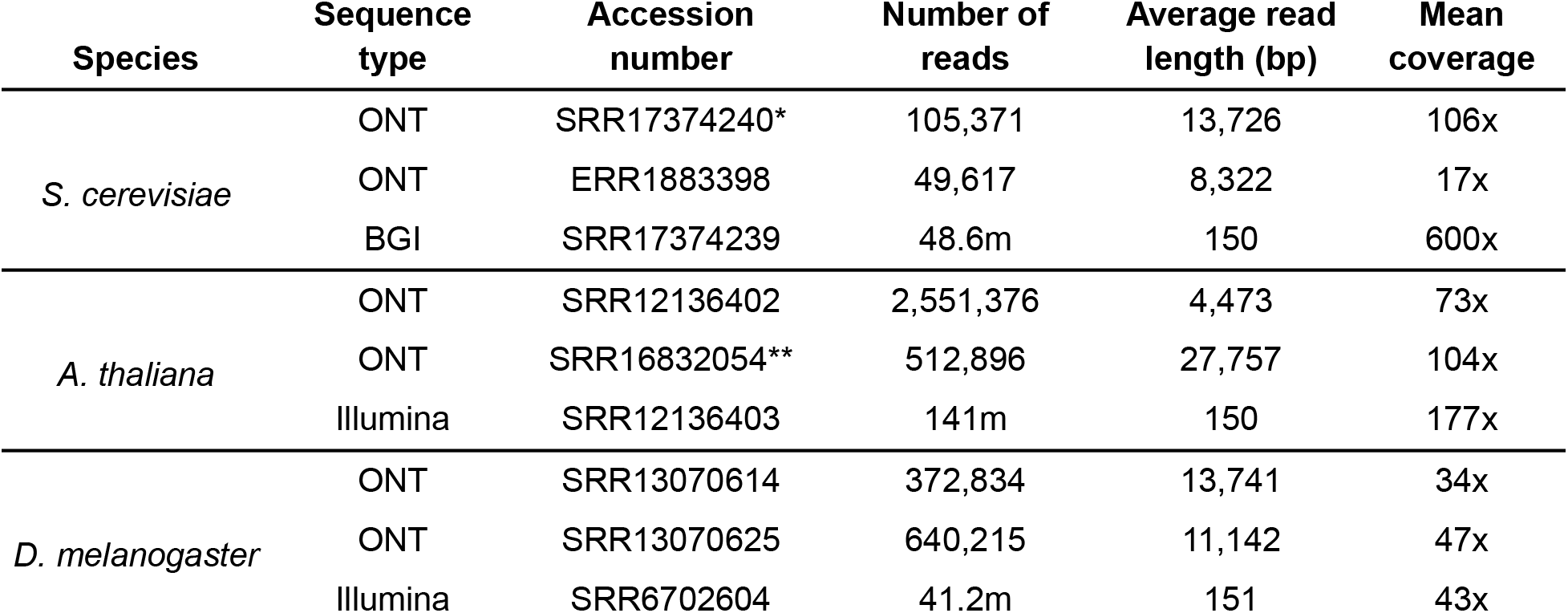
Experimental datasets used for performance evaluation. Seqtk was used to subsample *15% or **30% of the ONT reads for initial assembly.

**Table 3:** Simulated datasets used for performance evaluation.

| Species | Simulator | Number of reads | Average read length (bp) | Mean coverage |
| --- | --- | --- | --- | --- |
| *S. cerevisiae* | Badread (long reads) | 19,185 | 24,841 | 102x |
|  | ART (short reads) | 8.1m | 150 | 100x |
| *A. thaliana* | Badread (long reads) | 19,540 | 24,306 | 100x |
|  | ART (short reads) | 79.6m | 150 | 100x |
| *D. melanogaster* | Badread (long reads) | 16,268 | 24,162 | 98x |
|  | ART (short reads) | 94.8m | 150 | 100x |

### Polishing and Performance Assessment

FMLRC2 v0.1.6 using RopeBWT2 (r187; Li, 2014), HyPo v1.0.3 (Kundu et al., 2019), NextPolish v1.4.0 (Hu et al., 2020), ntEdit v1.3.5 (Warren et al., 2019), Pilon v.1.24 (Walker et al., 2014), POLCA v4.0.8 (Zimin & Salzberg, 2020), Polypolish v0.5.0 (Wick and Holt, 2022), Racon v1.5.0 (Vaser et al., 2017) and wtpoa (Ruan & Li, 2019) were used to polish nanopore-only bacterial and eukaryotic assemblies. Each polisher was run once on each assembly using the default parameters, unless otherwise specified. The resulting polished eukaryotic assemblies were then compared against their respective reference genomes using QUAST v5.0.2 (Gurevich et al., 2013). Bacterial assemblies were compared against the respective reference for simulated data, or in a pairwise fashion for experimental data as described in Wick and Holt (2022). Briefly, global alignments were computed between polished assemblies and the reference or species-matches assemblies for the simulated and experimental datasets, respectively. Total residual errors - or total pairwise distance - are equivalent to the edit distance, including mismatches and indels. Computational performance (CPU time and RAM usage) were determined using “/usr/bin/time -v”.

## Results

We evaluated FMLRC2 against seven other state-of-the-art assembly polishing methods using a combination of simulated and experimental long- and short-read datasets spanning a wide variety of bacterial species and three eukaryotes. Since the ground truth is known for the simulated datasets and reference lines of eukaryotes, we evaluated based on total residual errors among polishing results for simulated bacteria and all eukaryotic datasets (Figures 1 and 3, respectively). Table 4 shows the average residual errors per 100Kbp among simulated and experimental datasets from eukaryotes and the average CPU time and memory usage per polishing run. For experimental bacterial data, we use the total pairwise distance among technical replicates as an indicator of polishing accuracy (Figure 2). Polishing with FMLRC2 results in dramatically lower residual error and pairwise distance among bacterial datasets. Likewise, it produces the fewest residual errors among eukaryotic datasets, albeit not dramatically lower than the other best performing methods. However, FMLRC2 requires far less CPU time than the other methods, and 15X faster than the next best-performing method, NextPolish. The memory (RAM) usage is comparable to the other high-performing methods (except ntEdit and wtpoa, which have noticeably poor polishing accuracy).

**Table 4.**
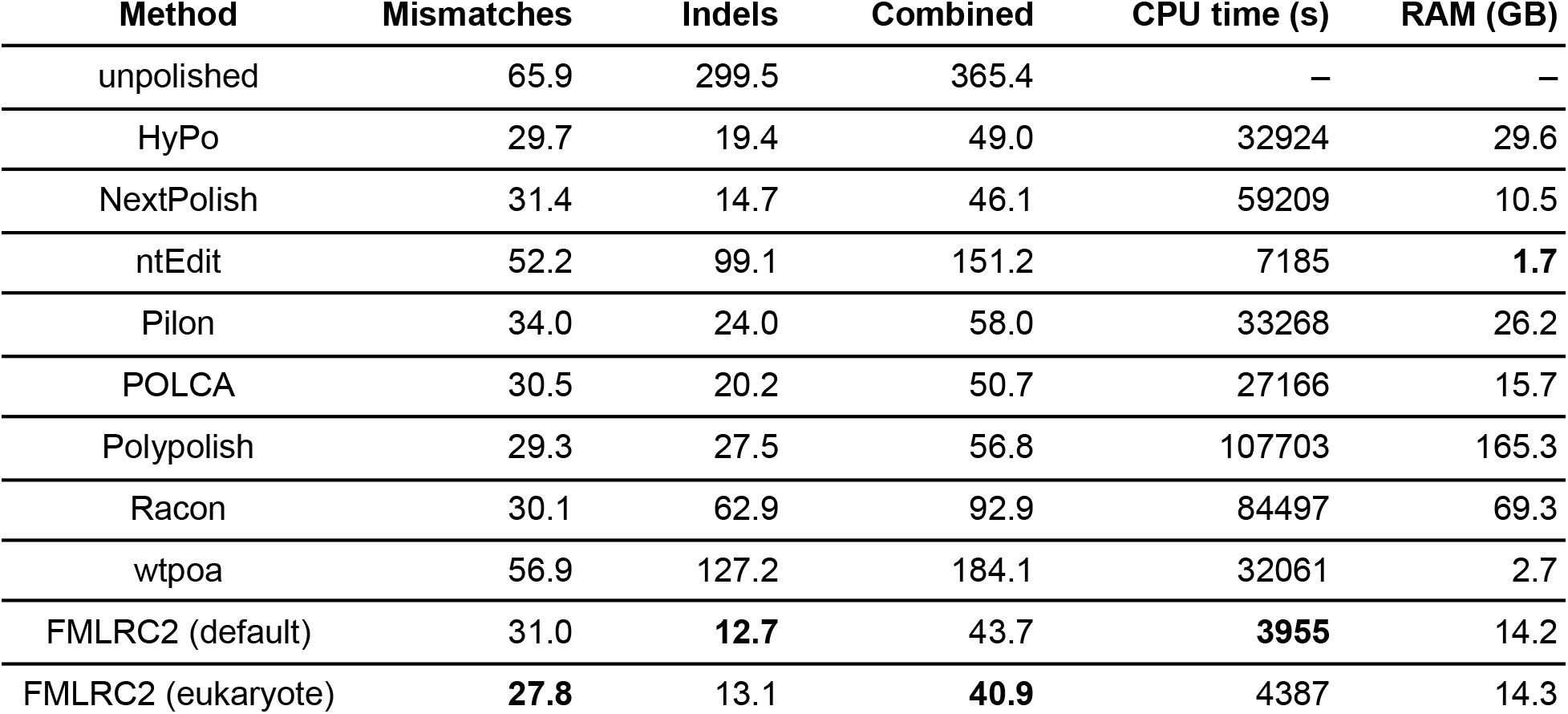
Performance of assembly polishers averaged over two experimental and one simulated dataset from each of the three species. Mismatches and indels represent the average residual errors per 100Kbp.

| Method | Mismatches | Indels | Combined | CPU time (s) | RAM (GB) |
| --- | --- | --- | --- | --- | --- |
| unpolished | 65.9 | 299.5 | 365.4 | – | – |
| HyPo | 29.7 | 19.4 | 49.0 | 32924 | 29.6 |
| NextPolish | 31.4 | 14.7 | 46.1 | 59209 | 10.5 |
| ntEdit | 52.2 | 99.1 | 151.2 | 7185 | **1.7** |
| Pilon | 34.0 | 24.0 | 58.0 | 33268 | 26.2 |
| POLCA | 30.5 | 20.2 | 50.7 | 27166 | 15.7 |
| Polypolish | 29.3 | 27.5 | 56.8 | 107703 | 165.3 |
| Racon | 30.1 | 62.9 | 92.9 | 84497 | 69.3 |
| wtpoa | 56.9 | 127.2 | 184.1 | 32061 | 2.7 |
| FMLRC2 (default) | 31.0 | **12.7** | 43.7 | **3955** | 14.2 |
| FMLRC2 (eukaryote) | **27.8** | 13.1 | **40.9** | 4387 | 14.3 |

**Figure 1.**
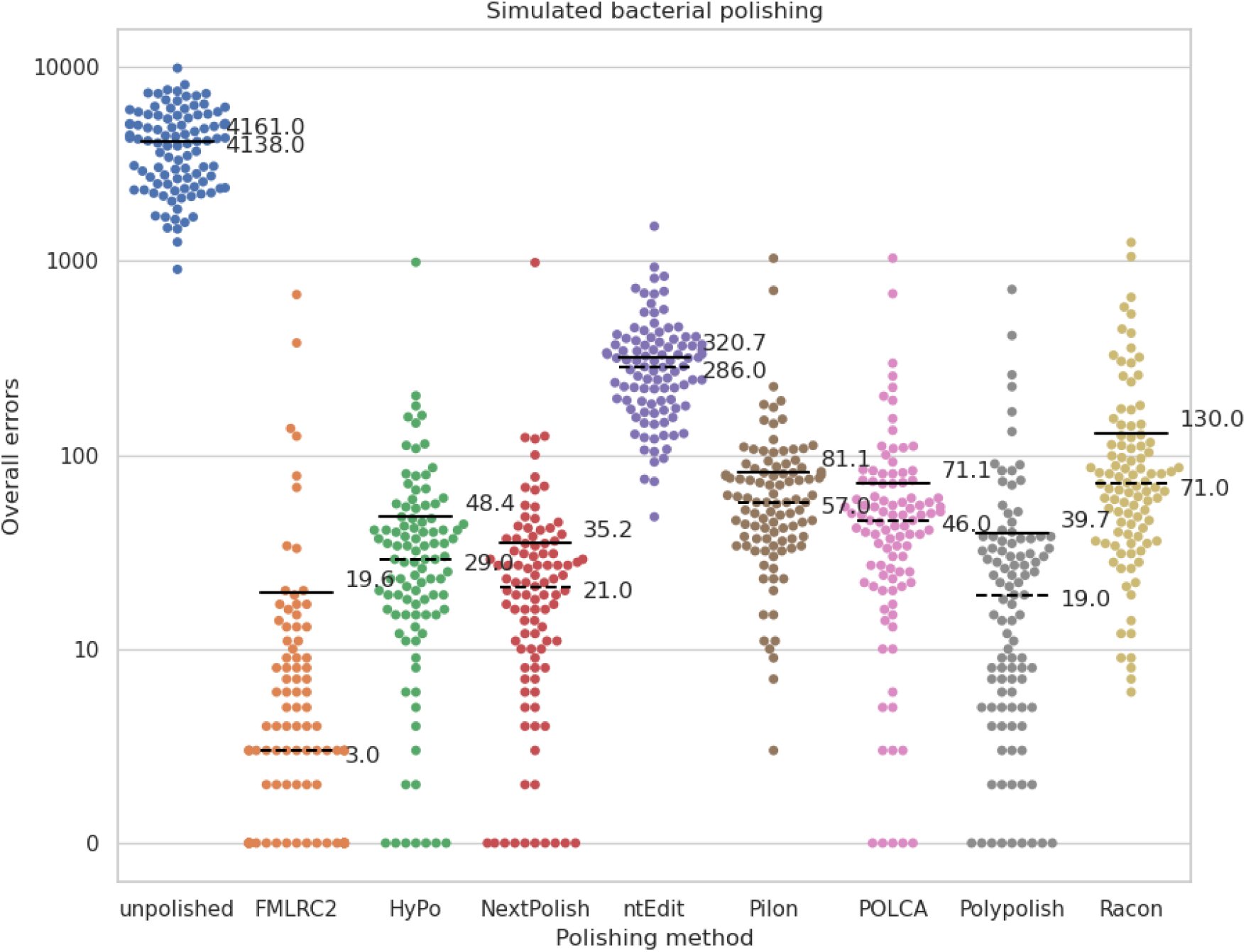
Overall error rates for simulated bacterial genomes. The solid line indicates the mean; dashed indicates the median.

**Figure 2.**
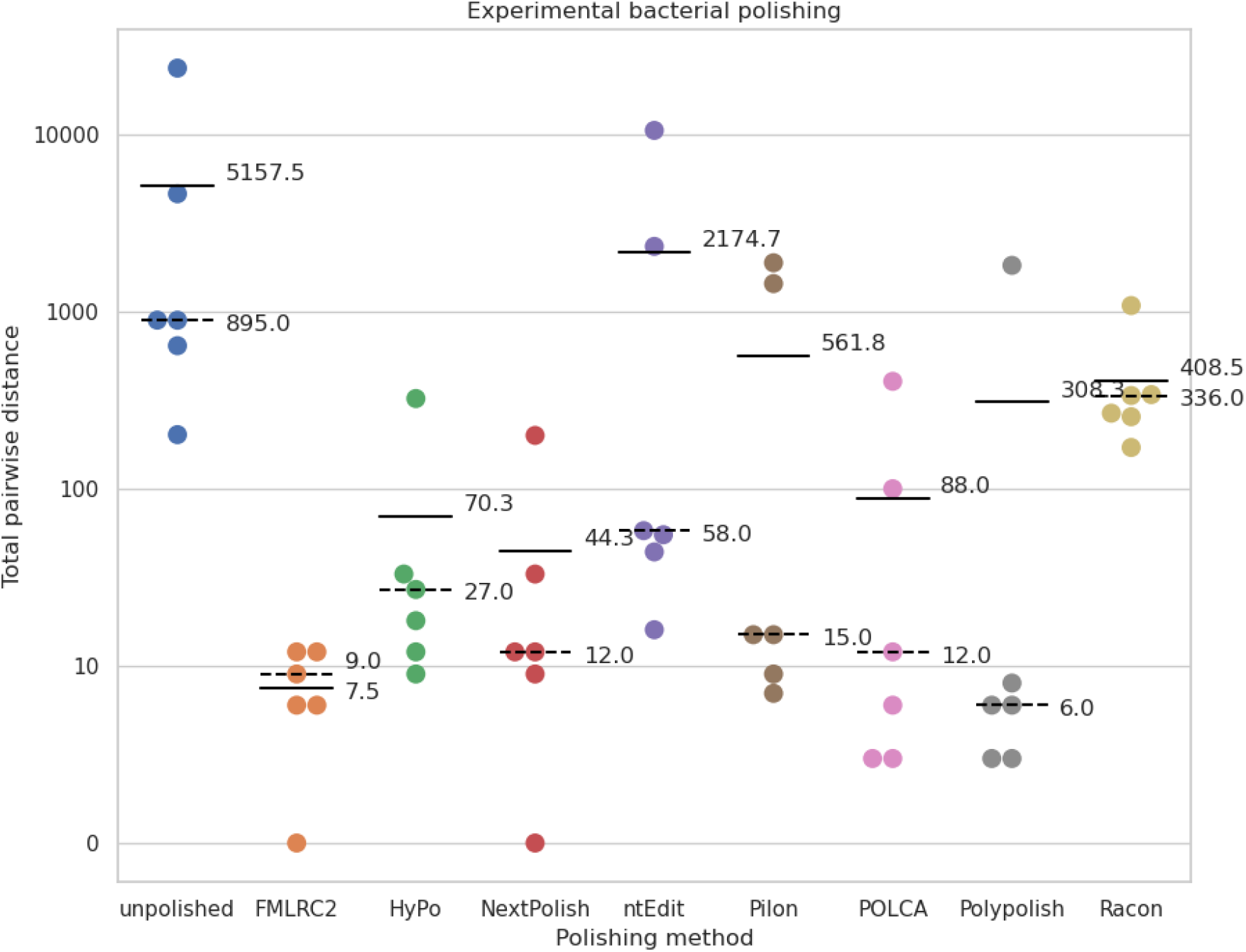
Sum of pairwise differences among replicates from experimental bacterial datasets. The solid line indicates the mean; dashed indicates the median.

**Figure 3.**
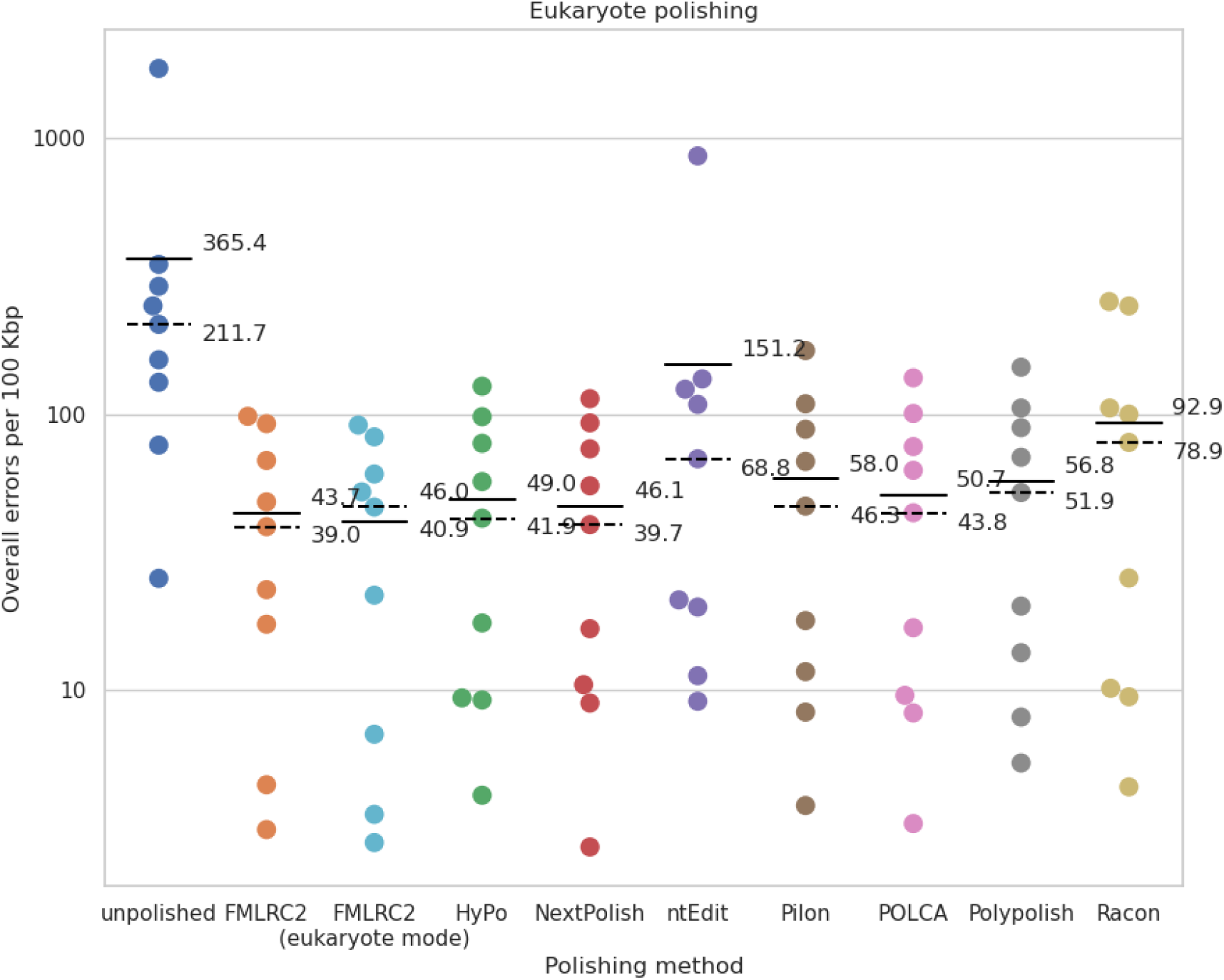
Residual errors after polishing for experimental and simulated eukaryotic datasets. The solid line indicates the mean; dashed indicates the median.

Results of polishing the *Drosophila melanogaster* datasets with FMLRC2 using its default parameters (--branch_factor 4, --cache_size 8, --k 21 59, --min_count 5, --min_frac 0.1) showed relatively poor performance correcting mismatch errors, especially among simulated datasets. We hypothesized that these errors occur in highly repetitive sequences (the likes of which do not exist in most bacterial genomes) when the representation of the true “version” of the repeat falls below the absolute or relative minimum count (min_count and min_frac, respectively). To address this, we evaluated all eukaryotic datasets using alternate parameter settings optimized for resolving these repetitive element problems (--k 21 59 80, --min_frac 0), dubbed “eukaryote mode”. With these settings, the average residual error across eukaryotic assemblies polished with FMLRC2 is further reduced at the cost of a ∼10% increase in CPU time (Figure 3, Table 4). Of note, however, FMLRC2 using the *default* settings still outperforms all other evaluated methods.

## Discussion and Conclusion

While the cost of 3rd-generation long read sequencing, including Oxford Nanopore Technologies, continues to decrease, and accuracy increases, hybrid multi-technology methods remain an efficient and effective approach for *de novo* genome assembly. Following on FMLRC’s proven performance as an error correction tool for raw reads, we demonstrate FMLRC2’s exceptional performance as a polishing tool for *de novo* nanopore-based assemblies in both bacteria and simple eukaryotes. FMLRC2 significantly outperforms the other tested polishing tools in reducing the residual assembly error, as illustrated using simulated and real ONT sequencing datasets. Assemblies polished with FMLRC2 have the fewest mean residual errors while FMLRC2 is also the fastest method - over an order of magnitude faster than the next most accurate tool.

